# Use of a predictor cue during a speech sound discrimination task in a *Cntnap2* knockout rat model of autism

**DOI:** 10.1101/2024.12.04.626861

**Authors:** Tracy M. Centanni, Logun P. K. Gunderson, Monica Parra

**Affiliations:** Department of Psychology, Texas Christian University, Fort Worth, TX 76129; Department of Speech, Language, and Hearing Sciences, University of Florida, Gainesville, FL 32610

**Keywords:** prediction, rodent, cerebellum, expectation, speech sound

## Abstract

Autism is a common neurodevelopmental disorder that despite its complex etiology, is marked by deficits in prediction that manifest in a variety of domains including social interactions, communication, and movement. The tendency of individuals with autism to focus on predictable schedules and interests that contain patterns and rules highlights the likely involvement of the cerebellum in this disorder. One candidate-autism gene is contact in associated protein 2 (*CNTNAP2*), and variants in this gene are associated with sensory deficits and anatomical differences. It is unknown, however, whether this gene directly impacts the brain’s ability to make and evaluate predictions about future events. The current study was designed to answer this question by training a genetic knockout rat on a rapid speech sound discrimination task. Rats with *Cntnap2* knockout (KO) and their littermate wildtype controls (WT) were trained on a validated rapid speech sound discrimination task that contained unpredictable and predictable targets. We found that although both genotype groups learned the task in both unpredictable and predictable conditions, the KO rats responded more often to distractors during training as well as to the target sound during the predictable testing conditions compared to the WT group. There were only minor effects of sex on performance and only in the unpredictable condition. The current results provide preliminary evidence that removal of this candidate-autism gene may interfere with the learning of unpredictable scenarios and enhance reliance on predictability. Future research is needed to probe the neural anatomy and function that drives this effect.

## Introduction

Autism is a common neurodevelopmental disorder that with a complex etiology but is notably marked by deficits in prediction that manifest in a variety of domains including social interactions, communication, and movement. Given the range of prevalent deficits, individuals with autism tend to struggle more often in academics, are less likely to achieve independence, and have poorer long-term life outcomes than neurotypical individuals [14]. More specifically, individuals with autism often exhibit deficits in tasks that require creating and evaluating upcoming events. For example, children with autism often fail to catch a bouncing ball, likely due to an inability to predict its trajectory [32]. The social deficits in autism, including theory of mind, are also possibly due to an inability to predict or anticipate how another person might be thinking or feeling [4]. This prediction deficit is thought to underlie the tendency of individuals with autism to prefer predictable schedules and focus on interests that contain patterns and rules [7, 27, 29, 45]. Deficits with prediction or creating expectations about upcoming events in a number of domains are likely linked with abnormalities in cerebellar circuits [17, 46]. Given that the cerebellum is critically involved in evaluating anticipated movements and assisting the cortex in processing of speech, it is not surprising that the cerebellum is also one of the most consistently reported regions of anatomical and functional differences in autism [17]. The goal of the current study was to evaluate the effect of an autism gene known to impact cerebellar development on the detection of predictable vs. unpredictable rapid auditory stimuli in a rat model.

With respect to language, abnormalities in the cerebellum likely impact the development of speech sound knowledge early in life, with effects that cascade throughout the lifespan. In young adults, processing predictable sentences, for example, increases activation in the right cerebellum, which connects to the left cerebral language areas [18]. If cerebellar dysfunction in autism leads to an array of prediction deficits, understanding the direct link between the brain and behavior is important for developing earlier diagnostic tools, perhaps even prior to the onset of theory of mind in childhood. One method for probing whether prediction deficits are ubiquitous across modality and present early in life is through the study of autism-candidate genes.

Variants in the autism-candidate gene contactin associated protein 2 (*CNTNAP2*) are associated with sensory deficits and anatomical differences [3]. The protein coded for by this gene, Caspr2, plays a role in axonal growth for mature myelinated neurons and, when deficient, leads to abnormal neuronal migration [6]. This gene is highly expressed in Purkinje cells, the main neuron type in the cerebellum, and variants in this gene are associated with reduced cerebellar grey matter in humans [47]. Mice with *Cntnap2* knockout exhibited significant differences not only in Purkinje cell morphology, but also increased spontaneous firing, increased excitability, and earlier onset latencies in response to somatosensory stimuli [25]. Abnormalities in development of auditory perception and reactivity in *Cntnap2* knockout rats [41] align with symptoms reported in humans with autism and suggests this gene should be one of interest in sensory processing studies.

Most of the prior research characterizing the *Cntnap2* knockout rodent (including mice and rats) has investigated social behaviors, activity levels, and motor coordination (for review, see [31]). While cognitive flexibility has been evaluated in other genetic models, and atypical behaviors were reported in *Cntnap2* knockout mice [31], this has not yet been fully evaluated in the rat model, which is capable of much more complex operant behaviors. During a pre-pulse inhibition (PPI) task, in which a cue indicates an upcoming startling event, there were mixed results on behaviors such that *Cntnap2* knockout mice showed an impaired startle response [5] while the effect in rats varied based on genotype and age [42]. Adult rats with homozygous *Cntnap2* knockout exhibited increasing startle response during a PPI paradigm [42], suggesting an impaired ability to anticipate upcoming events. It is unknown whether removal of this gene would also interfere with prediction during an operant task not containing aversive stimuli.

Given the inability to conduct causal genetic link studies in humans, animal model work is necessary to answer this question. The use of human speech sounds in rat models of human communication impairments is well-validated. Rats are capable of learning to discriminate human speech sounds based on the initial consonant [23, 36], or the middle vowel [34]. They can accomplish this task in the presence of background noise [44], with acoustic degradation [38], and at various speeds [11], while performing at levels that mimic human performance curves. Studies of auditory cortex encoding of speech sounds in rat models of autism have highlighted potential mechanisms for speech perception differences in humans, including altered patterns of cortical firing [20, 21]. Genetic rat models of other neurodevelopmental disorders, such as dyslexia, have led to the discovery of new neural deficits in a subgroup of children [9, 10, 13]. Thus, although rats do not process speech sounds as language, they are a useful model for understanding the impact of genetic manipulation on operant behavior and neural encoding of speech sounds as auditory cues. Thus, the current study utilized a *Cntnap2* knockout rat model to evaluate the impact of this gene on a rapid speech sound discrimination task that included unpredictable as well as predictable target presentations.

## Methods

### Animals

Subjects were 17 Long-Evans rats supplied by the Medical College of Wisconsin. Of these animals, 9 were Crispr/Cas9 *Cntnap2* knockouts (N = 2 female) and the remaining 8 were wildtype littermate controls (N = 3 female). During behavioral training, rats were housed on a reverse 24-hour light:dark cycle (lights on, 7:00 pm-7:00 am) with food restriction to ensure motivation during behavioral training but were not allowed to fall below 85% of their pre-deprivation body weight. Rats began training at three months of age. All animal protocols were approved by the Texas Christian University Institutional Animal Care and Use Committee.

### Behavioral Paradigm

The first portion of the task was identical to our prior work [8, 11, 12}. In brief, rats were trained to discriminate a target speech-sound (/dad/) from a randomized string of speech-sounds (/bad/, /gad/, /sad/, and /tad/) presented at varying speeds (2, 4, 5, 6.7, 10 and 20 syllables per second/sps). Training was conducted over two 30-min sessions per day (5 days/week) in an operant chamber placed into a double-walled, soundproofed booth (Vulintus Inc., Lafayette, CO). The chamber contained a pellet dispenser and an infrared-monitored nose poke on opposing walls. The first stage of training used autoshaping to teach the rat to place their nose in an infrared-monitored nose poke to associate the sound of the target (/dad/) with the receipt of a 45 mg sugar pellet (sucrose) reward. After receiving at least 100 pellets in a single session, they were then trained to hold their nose in the nose poke until the target sound /dad/ was played, at which time they should remove their nose to receive the sugar pellet reward. Progression through training stages was dictated by previously demonstrated criteria using the d’ value of each training session as an indicator of discrimination ability (**Table 1**). The d’ value is defined as the Z-score of the distractor response rate subtracted from the z-score of the target response rate. After 10 nonconsecutive sessions with a d’ ≥ 1.50, a series of eight training stages was used to introduce each of the four distractors and increasing levels of randomization. After completion of stage 11, the rats were trained to discriminate the target sound from the distractors at progressively increasing speeds up to 20 sps. Next, they were placed into a new training stage, where the distractor sound /bad/ was converted to a predictor sound such that whenever /bad/ was played, the target sound always followed [28]. After achieving two nonconsecutive sessions with a d’ ≥ 1.50, rats completed a final testing stage over 10 sessions, during which the predictor was played in 40% of initiated trials, with the target appearing at random without the predictor in the remaining 60% of trials. Response rates and reaction times to each stimulus were recorded as dependent variables for analysis.

**Table 1.** Description of speech-discrimination training/testing stages and thresholds.

| Stages | Description | Thresholds |
| --- | --- | --- |
| 1 | Autoshaping | One session with 100 pellets |
| 2 | Train to hold in nose poke until target sound /**dad**/ is presented | 10 nonconsecutive sessions with $\geq 30$ pellets and $d' > 1.5$ |
| 3-6 | Each stage introduces one of the four distractor sounds, /**tad**/, /**gad**/, /**sad**/, /**bad**/, with the target | Two sessions with $\geq 30$ pellets and $d' > 1.5$ |
| 7-11 | Each stage increases the number of distractors presented in a given block of trials, until all five sounds are presented | Two sessions with $\geq 30$ pellets and $d' > 1.5$ |
| 12 | Introducing faster presentation rate of 4 sps | Eight sessions with $\geq 30$ pellets and $d' > 1.5$ |
| 13-16 | Introduce rest of increased presentation rates of 5, 6.7, 10, and 20 sps | Four sessions with any $d'$ |
| 17 | Randomized presentation rates | 10 sessions with any $d'$ |
| 18 | Convert distractor /**bad**/ to a predictor, where /**dad**/ always follows when presented | Two sessions with $\geq 30$ pellets and $d' > 1.5$ |
| 19 | Testing stage for prediction, 40% of initiated blocks of trials will have predictor /**bad**/ presented | 10 sessions with any $d'$ |

### Statistical Plan

All analyses were performed using custom code in Matlab (version R2021b). Response rate was defined as the total number of responses to each sound at each presentation rate with respect to the total number of presentations. Reaction time was defined as the mean time to respond to a stimulus in milliseconds (ms). Repeated measures analyses of variance (ANOVA) were utilized to assess the effect of the predictor (present vs. absent), presented sound, and presentation rate on response rates and reaction times. Any significant main effects and interactions were explored with post-hoc *t-*tests using the Bonferroni adjustment to correct for multiple comparisons as needed.

## Results

### Knockout rats were affected by randomized distractors during training

Both groups of animals progressed through training at the same pace such that the number of days to hit criterion in each training stage was equivalent across genotype groups (one-tailed unpaired t-tests; *ps* > 0.10). When randomization was first introduced in training (stage 11; [11, 12]), the knockout rats responded significantly more to the distractors (13.66 ± 1.41%) compared to the wildtype controls (9.78 ± 1.57%; one-tailed t-test, *t* (15) = 1.97, *p* = 0.03), but responded just as frequently to the target sound (KO: 64.89 ± 5.16 vs. WT: 61.38 ± 3.53%; one-tailed t-test, *t* (15) = 0.58, *p* = 0.28; **Fig 1**). As training progressed, knockout rats continued to respond significantly more often to the distractors at the slowest speed compared to the wildtype controls (*ps* < 0.03). To ensure this was not due to an anxiety behavior or a motor deficit, we also evaluated reaction times during training to all stimuli. Across genotypes, there were no significant differences between reaction times to the distractors (two-tailed unpaired t-test, *t* (15) = 0.33, *p* = 0.74) or to the target sound (two-tailed unpaired t-tests, *t* (15) = 1.00, *p* = 0.33). Within the wildtype control group, rats generally responded more quickly when false alarming to distractors vs. responding to the targets (two-tailed paired t-test; *t* (7) = 2.80, *p* = 0.026). Within the knockout group, there was a trend in the same direction (two-tailed paired t-test; *t* (8) = 2.21, *p* = 0.058).

**Fig 1.**
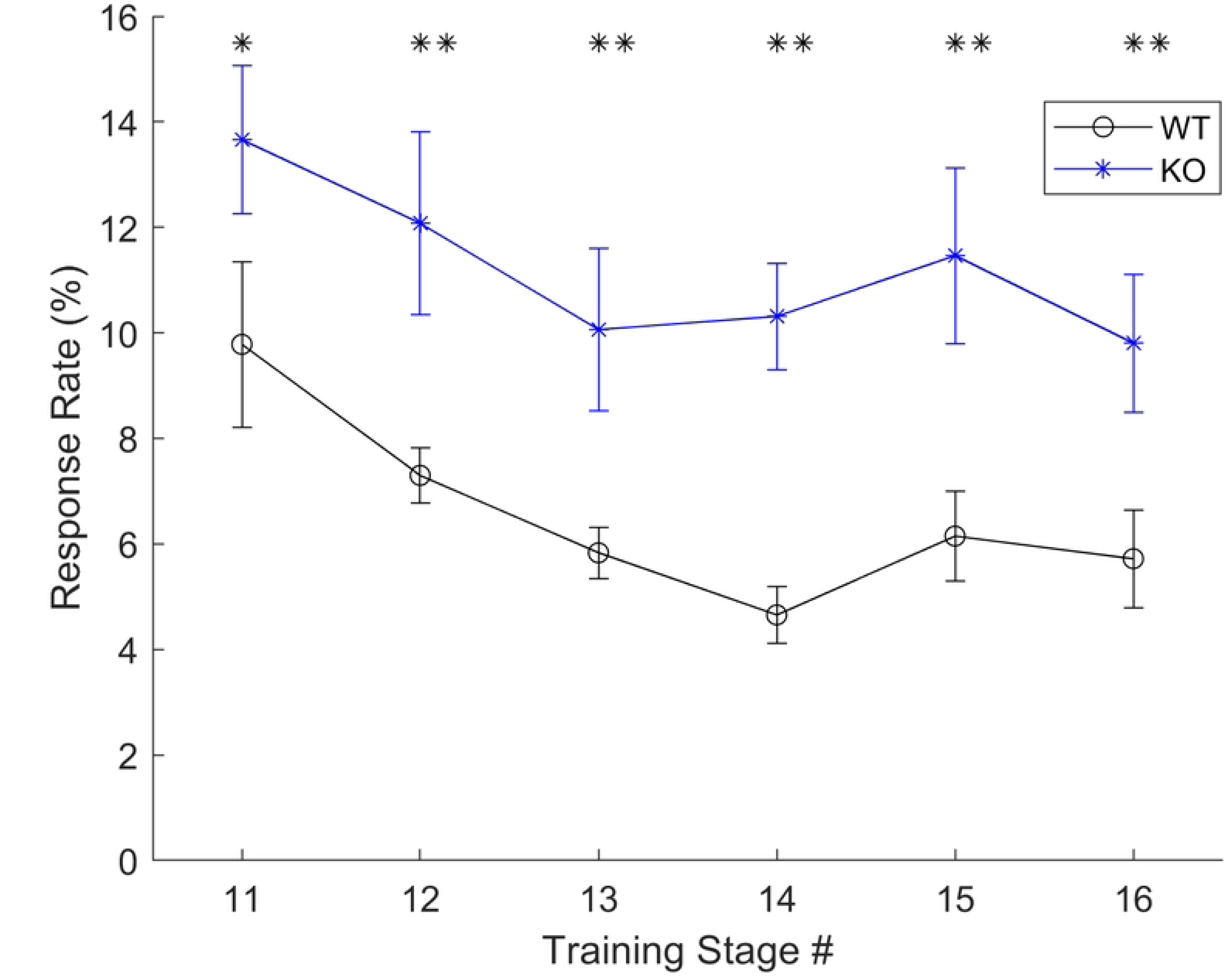
False alarm rates were higher in KO rats during early training. In stages 11-16 of training (after rats learned the mechanism of response and reliably responded to the target sound), rats were gradually exposed to more complex unpredictable sequences and faster presentation rates [12]. During these stages of training, knockout rats responded more frequently to the distractors (increased false alarms) compared to the wildtype controls. * p < 0.05, ** p < 0.01

### Knockout rats can efficiently utilize a predictor cue

As expected based on our prior work [11, 12, 28], wildtype controls learned the unpredictable rapid speech sound discrimination task and learned to utilize the predictor cue to anticipate the target sound /dad/ (**Fig 2**; black lines). The KO rats similarly learned both versions of the task, exhibiting comparably high hit rates in both predictable and unpredictable conditions during final testing (**Fig 2**; blue lines). There were significant main effects of genotype (F (1, 1008) = 24.5, *p* < 0.0001), presentation rate (F (5, 1008) = 33.77, *p* < 0.0001), predictability (F (1, 1008) = 48.44, *p* < 0.0001), and sound (F (4, 1008) = 232.14, *p* < 0.0001). Within each genotype group, we evaluated the effect of the predictor on hit rates and false alarm rates. In the WT group, hit rates to the target sound were higher when the predictor was present at 2 sps (two-tailed, paired t-tests; *t* (7) = 3.6, *p* = 0.009) but hit rates were lower when the predictor was present at 6.7 sps (*t* (7) = 2.51, *p* = 0.04). This reduced hit rate in the context of the predictor could be due to the learned relationship between predictor and target leading to increased anticipatory responses to the target [28]. False alarm rates in the WT group were, however, higher in the unpredictable condition at the four slower speeds compared to in the presence of the predictor (*ps* < 0.038). In the KO group, there were no differences in hit rates in the predictable vs. unpredictable conditions (*ps* > 0.17). There were differences in false alarm rates in the KO group such that rats exhibited higher false alarm rates at every speed in the unpredictable condition (*ps* < 0.05).

**Fig 2.**
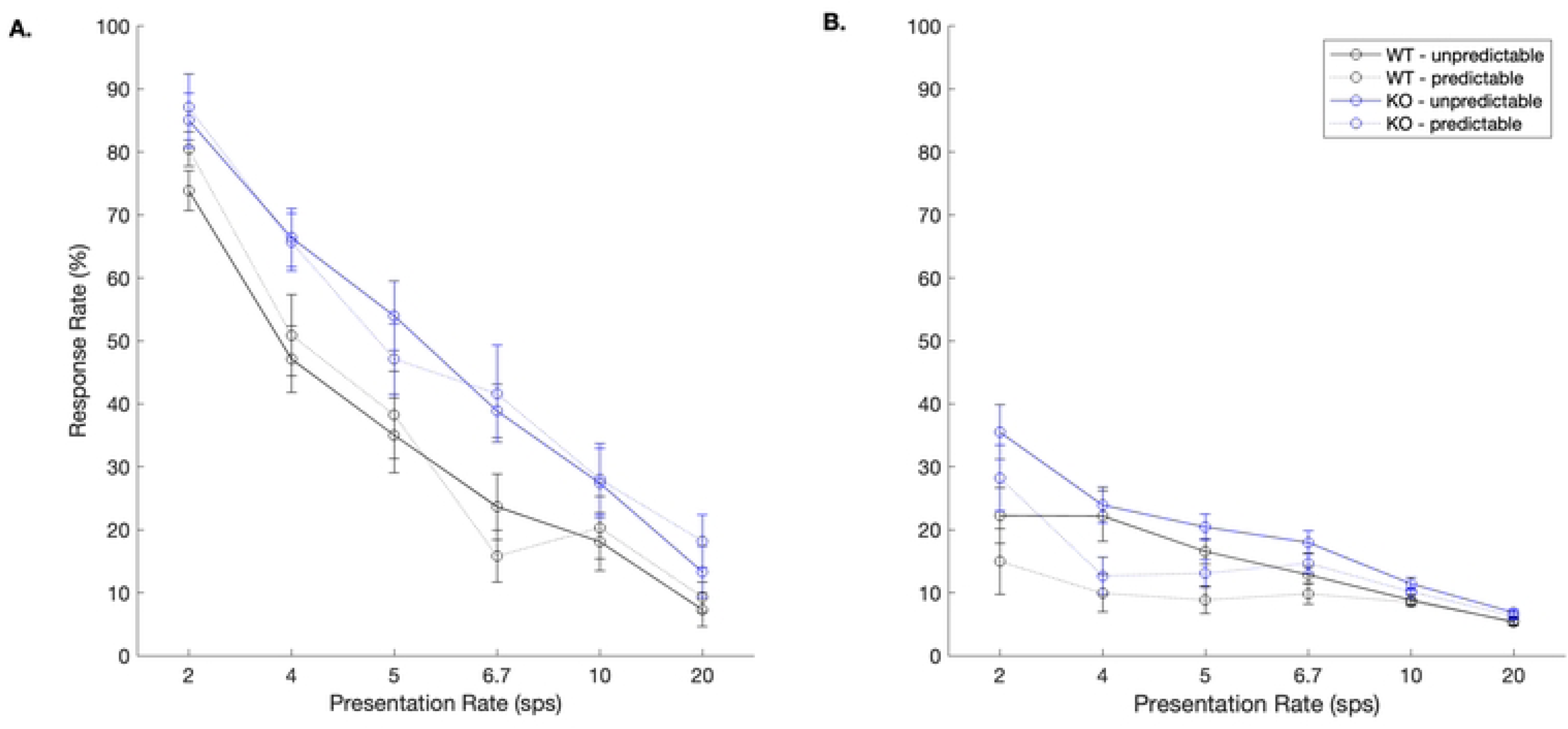
Rats can learn to identify a target sound regardless of genotype. **A.** Hit rates to the target /dad/. **B.** During unpredictable trials, summed false alarm rates to the three distractors were significantly below hit rates until 20 sps (one-tailed paired t-tests; WT *ps* < 0.039, KO *ps* < 0.01), as has been shown in our prior work [12]. During predictable trials, KO rats responded more to the target than the distractors at all speeds (*ps* < 0.011) and WT rats exhibited this pattern at 4 of the 6 speeds (all except 6.7 and 20 sps; *ps* < 0.02).

Interestingly, early anticipatory response rates to the predictor sound /bad/ were significantly higher in the KO group at the two slowest and the two fastest speeds (one-tailed t-tests, *ps <* 0.04; **Fig 3**). This increased anticipatory response, however, did not impede accuracy. The KO animals responded significantly more frequently to the target sound /dad/ than the WT rats at 4 sps (KO: 65.70 ± 4.59 vs. WT: 50.91 ± 6.43%; one-tailed t-test, *t* (15) = 2.03, *p* = 0.03), 6.7 sps (KO: 41.62 ± 7.73 vs. WT: 15.81 ± 4.13%; *t* (15) = 3.01, *p* = 0.004), and 20 sps (KO: 18.20 ± 4.20 vs. WT: 9.30 ± 2.37; *t* (15) = 1.89, *p* = 0.039). This pattern of results suggest that the KO group was more efficient at utilizing the predictor cue to minimize false alarms and more efficiently respond to the target sound.

**Fig 3.**
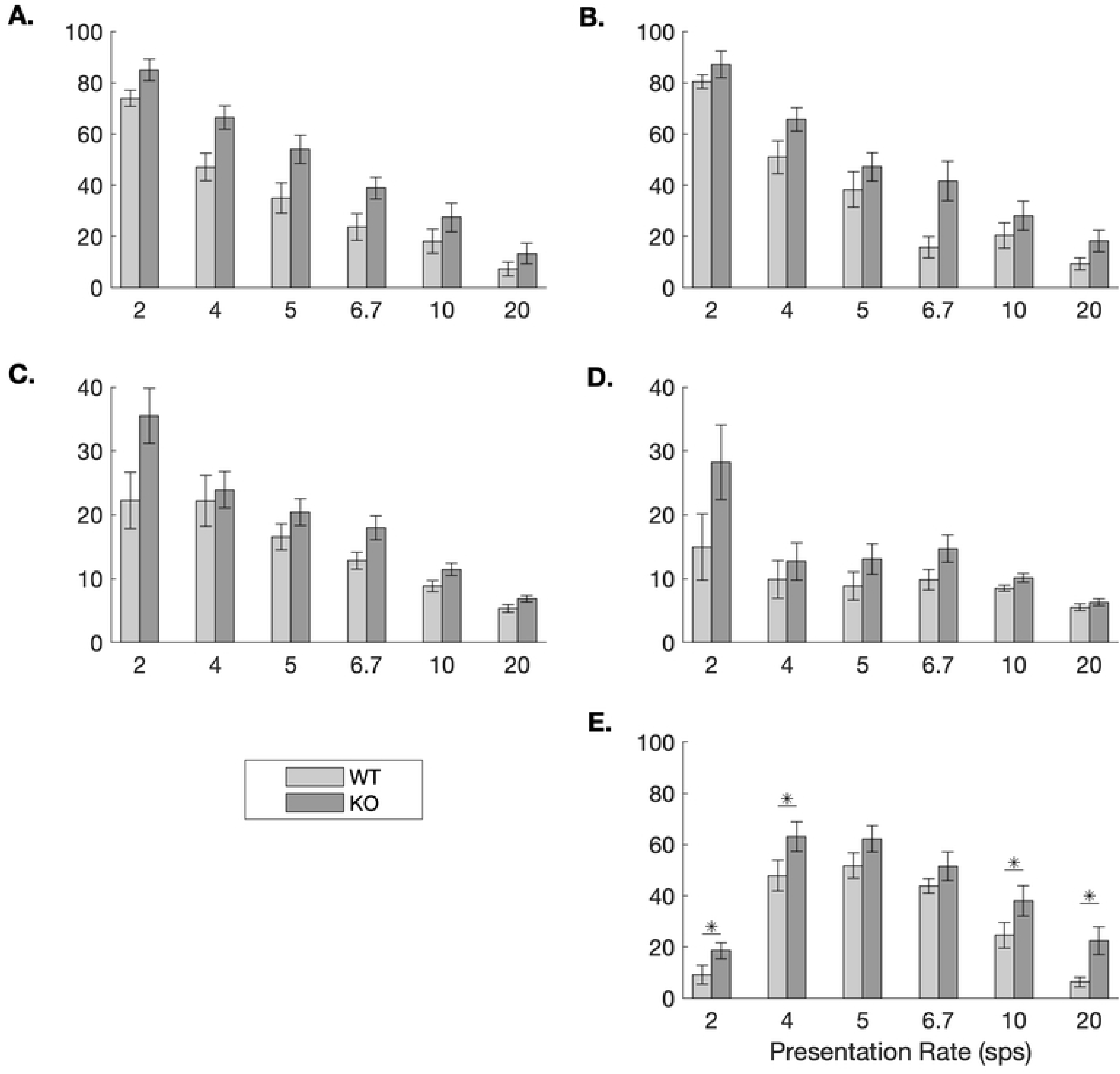
Response rates by condition and genotype. During the testing stages, there were no differences in response rates to the target sound /dad/ across genotypes in the unpredictable (**A.**) or predictable (**B.**) conditions. While there was no effect of predictability on false alarm rates in the WT rats, the KO rats exhibited higher false alarm rates during unpredictable trials (**C.**) compared to predictable trials (**D.**) During predictable trials, the response to the predictor sound /bad/ was higher in the KO group compared to the WT group at four of the six speeds (**E.**).

With respect to reaction times, in the unpredictable condition, WT rats responded significantly faster to the target /dad/ compared to the KO rats at three speeds: 5 sps (unpaired, two-tailed t-test, *t* (15) = 2.13, *p* = 0.05), 6.67 sps (*t* (15) = 2.34, *p* = 0.03), and 20 sps (*t* (15) = 2.32, *p* = 0.03; **Fig 4**). The same general pattern was observed during the prediction condition. During these trials, WT rats were significantly faster at responding to the target /dad/ at 6.7 sps (*t* (15) = 2.88, *p* = 0.01) and 20 sps (*t* (15) = 3.99, *p* = 0.001).

**Fig 4.**
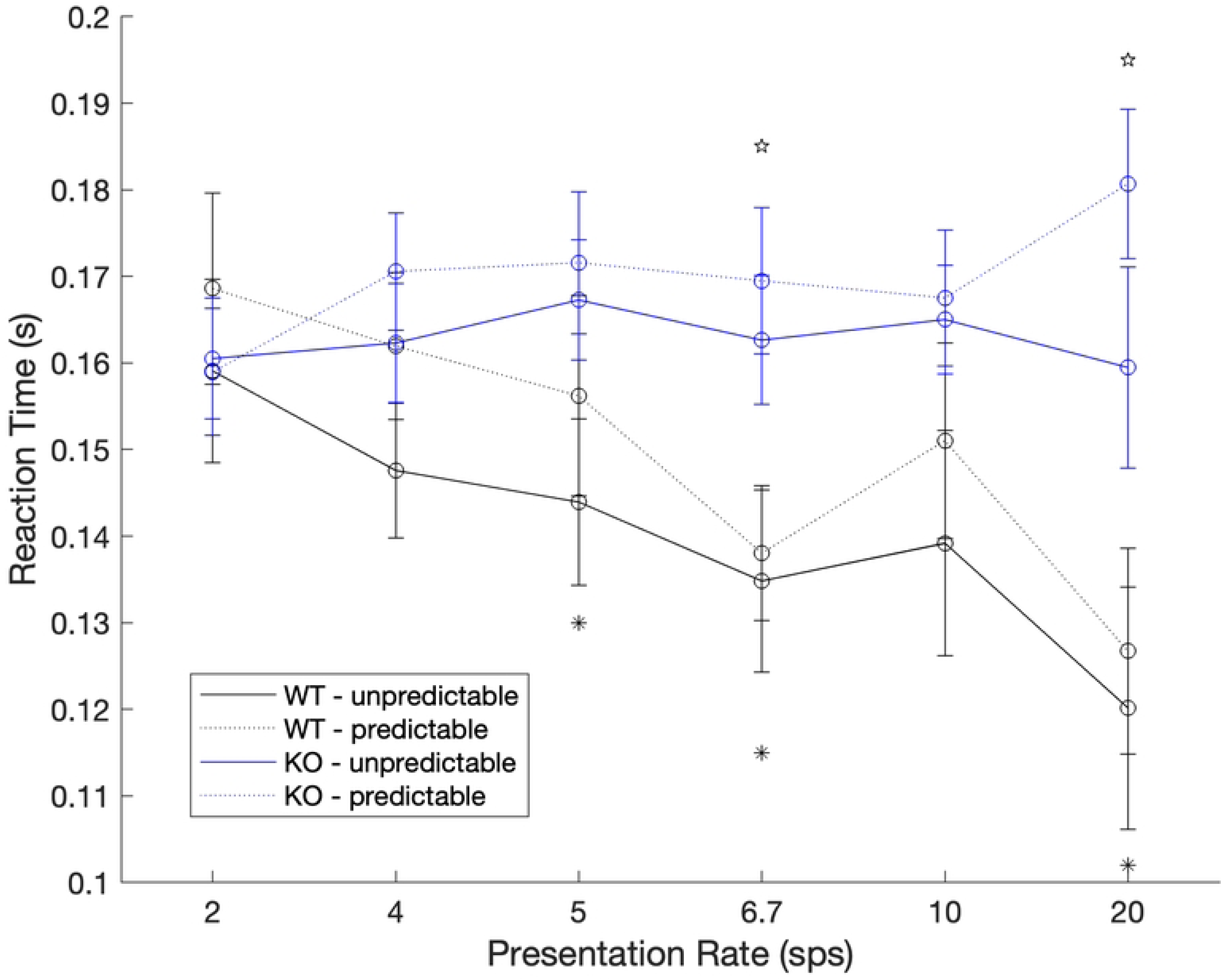
WT rats respond faster to the target than KO rats. During the unpredictable trials, WT rats responded faster to the target sound than KO rats at three speeds (* p < 0.05). During predictable trials, a similar pattern was observed, where WT rats responded faster than KO rats at two of the six speeds (∂ p < 0.05).

### Preliminary evidence for sex differences

Given the small numbers of female rats in the sample (N = 3 WT, N = 2 KO), we conducted an exploratory analysis on possible sex differences on response rates. First, we evaluated response rates in the unpredictable condition. In the WT group, female rats responded significantly more to the target /dad/ at 10 sps (29.68 ± 6.62%) compared to males (11.15 ± 3.74%; unpaired, two-tailed t-test, *t* (6) = 3.11, *p* = 0.02). In the KO group, female rats responded significantly more to the target /dad/ at 20 sps (27.88 ± 4.08%) compared to males (9.12 ± 3.80%; unpaired, two-tailed t-test, *t* (7) = 2.67, *p* = 0.03). In the predictable condition, there were no sex differences in response rates to the target /dad/ in the WT group (*ps* > 0.22) or in the KO group (*ps* > 0.16). Similarly, there were no sex differences in response rates to the predictor /bad/ in the WT group (*ps* > 0.19) or in the KO group (*ps* > 0.58).

## Discussion

The goal of the current study was to evaluate the effect of *Cntnap2* knockout on the ability of rats to learn and utilize a predictive cue during a rapid auditory discrimination task. Although both knockout and wildtype rats were able to learn the task and perform with comparable accuracy during unpredictable trials, the knockout rats exhibited significantly higher false alarm rates during training and higher hit rates to the target during testing when the predictor cue was available compared to their wildtype counterparts. The current results provide preliminary evidence that removal of this candidate-autism gene creates an animal that is more likely to respond to a distractor in unpredictable conditions and performs more accurately in predictable conditions. Future research is needed to probe the neural anatomy and function that drives this effect and consider translation of these findings into humans.

### Cntnap2 KO rats utilize a predictor cue more effectively

In the current study, we observed evidence of improved use of a predictor cue in rats with *Cntnap2* KO compared to WT controls. Our KO rats were more accurate during predictable trials at 4, 6.7, and 20 sps speeds compared to their WT counterparts, with the largest effect at 6.7 sps. While *Cntnap2* KO rats exhibited more false alarms than WT rats during initial training on random sequences, this effect disappeared by the testing phases, when there was no main effect of predictability. One possible explanation for the lack of a deficit during unpredictable sequences in our study is that the prediction deficits associated with abnormal *Cntnap2* expression are not universal and may be restricted to certain types of behaviors. For example, prior studies in rodent models have reported social deficits following *Cntnap2* knockout [40] and social deficits are associated with this gene in humans [15]. Since we did not use conspecific vocalizations and human speech is not meaningful to rats, we cannot speculate about whether this gene impacts speech sound perception in humans.

Our result also contradicts prior findings that rats with homozygous *Cntnap2* knockout exhibited increasing startle response during a pre-pulse (PPI) paradigm with age [42], suggesting an inability to anticipate upcoming events even when a predictor cue is available. This apparent contradiction may be explained by the fact that PPI tasks are often considered classical conditioning tasks, which may utilize different neural circuitry than is used for operant behaviors, as in the current task. In addition, the startling stimulus in a PPI task is often aversive (hence the startle response) whereas the target stimulus in our task was associated with a positive reinforcer. Thus, a consideration when interpreting our findings is the influence of this gene on specific neural circuitry involved in generating and evaluating expectations for aversive versus appetitive outcomes.

### Neural mechanisms for prediction

The cerebellum plays a fundamental role not only in evaluating predictions but also in adjusting to errors. For example, during walking, the cerebellum receives a copy of the movement plan and then utilizes somatosensory feedback to evaluate the execution of the plan. If an error occurs, the cerebellum registers the error and sends the signal to the cortex for a correction to be issued [30]. The removal of *Cntnap2* from the rat model in our study may have selectively interfered with part of this function. During the unpredictable condition of our task, there is no known plan of action or method available to anticipate the location of the target sound. Therefore, in this condition the auditory cortex may drive the behavior without the need for cerebellar involvement, and prior work has demonstrated that auditory cortex activation is sufficient to predict behavior in the rat during isolated and randomized speech sound discrimination tasks [11, 19]. During the predictable condition however, the cerebellum may be using the predictor as the anticipated event, comparing each sound against the predictor. It is unclear, however, how the removal of *Cntnap2* would lead to this enhanced ability to utilize a predictive cue.

*CNTNAP2* is expressed in the nodes of Ranvier, the spaces between myelin sheath on the axon [35]) and is involved in stabilizing the conduction of an action potential [49]. Removal of *Cntnap2* in a mouse led to significant alterations in action potentials in several cortical regions such that there was an increase in neurotransmitter release and excitatory post-synaptic potential amplitudes as well as an increase in repetitive motor behaviors [43]. Of note, though, these studies were conducted in cortical pyramidal neurons, and it is possible that knockout of *Cntnap2* impacts cerebellar Purkinje neurons differently. Future research is needed to probe the relationship between altered action potential transmission and reliance on predictability in behavioral tasks. The authors did suggest that this effect on action potential transmission may lead to abnormalities in connectivity and cortical circuit functions [49], which may provide an additional clue about the role of *CNTNAP2* in humans.

The effect of *CNTNAP2* variation on autism may be best understood at the circuit level, as supported also by its association with epilepsy and seizure activity in both humans [26, 39] and rodent models [33, 48]. Anatomical [16] and functional [50] connectivity are both abnormal in humans with *CNTNAP2* risk variants. Mice with *Cntnap2* risk alleles also exhibit abnormal neural network activity [33]. Interestingly, although *CNTNAP2* is expressed in Purkinje neurons in the cerebellum, this gene has higher expression levels in frontal and temporal lobes in humans [1]. This cortical bias in gene expression is not observed in the rodent brain, suggesting that the impact of *CNTNAP2* variation in humans may be meaningfully different than in a rodent model. If the circuit connecting cortical language and cognitive regions to the cerebellum is functioning at increased excitability [43], it may better explain not only the enhanced reliance on predictive cues compared to the rodent model, but also the inability of humans with autism to generate expectations in cortex and pass those to cerebellum for monitoring.

### Limitations

There are three main limitations in the current study. First, we did not collect data about brain function or anatomy. Given the known associations between *Cntnap2* and cerebellar abnormalities in rodent models [5, 25] as well as in humans [16, 47], future work is needed to determine whether the behavioral results we present here are attributed to these published differences or another circuit. Second, we were not adequately powered for statistical analyses on possible sex differences. Though we designed the study to support sex difference analyses, two animals experienced seizures during the study period, which is a known risk associated with *Cntnap2* knockout [33, 48], and another failed to learn the task. Given that autism presents differently in males compared to females, future study of the biological mechanisms of autism should include well-powered comparisons of sex. Finally, human speech sounds are not ecologically relevant to rats. These stimuli have been utilized in many prior preclinical studies of human developmental disorders, including dyslexia [10, 8], autism [20, 21], fragile X [22], Rett syndrome [2, 24], and others [36, 37, 38, 44]. Although these prior studies have demonstrated the relevance and utility of using human speech sounds in rodent model work, we cannot say with certainty that knockout of *Cntnap2* creates the same speech sound perception impairments observed in humans with autism. Thus, these results should be considered with this caveat in mind.

## Conclusion

The present study utilized a rat model to determine the impact of *Cntnap2* knockout on the ability to discriminate speech sounds in a predictable vs unpredictable context. We saw that *Cntnap2* knockout rats did respond to distractor stimuli early in training with fully randomized presentation of the sounds but outperformed their wildtype counterparts on test trials where the target sound was preceded by a predictive cue. These findings are in line with prior work showing impaired auditory discrimination to randomly presented stimuli and, perhaps more importantly, they also present novel evidence that rats with a knockout of this autism-associated gene can utilize a predictive cue more effectively than wild-type rats, Future work is needed to probe the specific neurological mechanisms that underlie these behavioral differences and translate these observations into the human population.

## Acknowledgments

This work was partially funded by 1R15HD103479-01A1 to TMC. The authors would like to thank Mya Conley, Alyssa Gutierrez, Sarah Grace White, and Kate Moorman-Wolfe for their assistance with behavioral training and handling of research animals.

